# Disentangling genetic feature selection and aggregation in transcriptome-wide association studies

**DOI:** 10.1101/2020.11.19.390617

**Authors:** Chen Cao, Devin Kwok, Qing Li, Jingni He, Xingyi Guo, Qingrun Zhang, Quan Long

## Abstract

The success of transcriptome-wide association studies (TWAS) has led to substantial research towards improving its core component of genetically regulated expression (GReX). GReX links expression information with phenotype by serving as both the outcome of genotype-based expression models and the predictor for downstream association testing. In this work, we demonstrate that current linear models of GReX inadvertently combine two separable steps of machine learning - feature selection and aggregation - which can be independently replaced to improve overall power. We show that the monolithic approach of GReX limits the adaptability of TWAS methodology and practice, especially given low expression heritability.

---

Transcriptome-wide association studies (TWAS) have successfully identified many associations between genes and complex traits^1–5^ and have triggered extensive methodological research^6–12^. The key concept behind TWAS is Genetically Regulated eXpression, or GReX, which is the component of gene expression attributed to genetic regulators. A typical TWAS procedure involves training a linear model, such as ElasticNet^6^ or Bayesian regression^7,8,13^, to estimate GReX as a weighted linear combination of regulatory DNA elements. The predicted GReX is then associated to phenotype in a separate association mapping dataset in which expression data is unavailable. The importance of GReX is evidenced by the number of publications which either seek to improve its prediction accuracy or expand its applications. Researchers have refined the original ElasticNet- and Bayesian-based models of PrediXcan^6^ and BSLMM^13^ by integrating multiple tissues^14–16^, adding trans-eQTLs^11^, and incorporating improved Bayesian methods^8^. GReX counterparts have also been developed for LD-score^17^, polygenic risk score^18^, and fine-mapping^12^.

However, recent efforts in improving GReX may have overlooked its primary purpose, which is not to predict expressions but to gather relevant genetic variants for association mapping^6^. This viewpoint is supported in practice by the low predictive accuracy of GReX models, which generally have an R^2^ value of 5% −10% for the topmost candidates due to low expression heritability^2,6,19,20^. Properly speaking, GReX is a weighted linear combination of genotypes which are selected via the objective of predicting expression data, and as such, these linear combinations do not necessarily represent true biological causes of gene expression. This subtle but important distinction suggests that understanding the statistical roles underlying GReX may yield greater benefits over simply optimizing its prediction accuracy. From the perspective of machine learning on high-dimensional data, GReX combines feature selection, which reduces dimensionality by removing irrelevant features (genetic variants), and feature aggregation, which calculates aggregate statistics from selected features (variants) to maximize statistical power (**Fig. 1.a**). The process of training the expression model is equivalent to using a linear model to select variants, and the process of associating predicted expressions to phenotype is equivalent to using the same linear model to aggregate variants (**Fig. 1.b**). Given this interpretation, we investigate whether feature selection and feature aggregation can be separated into two different methods, and if so, are there methods that perform better than the GReX linear model in either step (**Fig. 1.c**)? As our analyses of real and simulated data show, the answer to both questions is yes: it is not only possible to conduct feature selection and aggregation separately, we also propose a simple combination of methods that can outperform the use of GReX alone.

**Figure 1:**
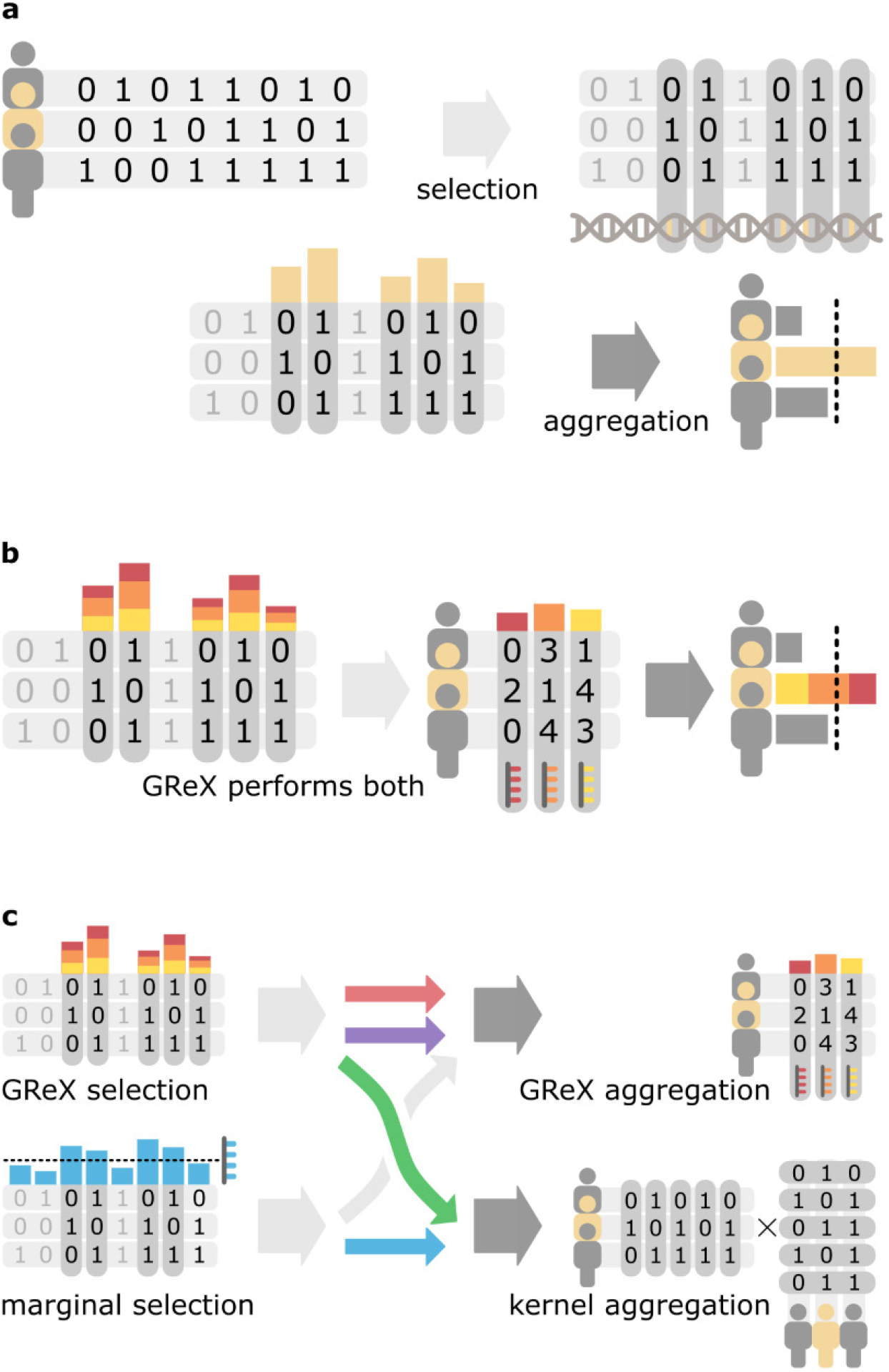
Function of GReX in terms of feature selection and feature aggregation. (**a**) Feature selection and feature aggregation are two typical steps in the statistical analysis of high-dimensional data. (**b**) The current practice of TWAS combines feature selection and feature aggregation into a single multiple linear regression model for estimating genetically regulated expression (GReX). (**c**) Separating these two fundamentally different steps allows a larger combination of methods to be applied to TWAS, providing greater flexibility in practice and potentially increased power. This study quantifies the performance of the four combinations illustrated with colored arrows.

This work extends the findings of two recent publications which replace GReX-based feature aggregation in TWAS with kernel-based methods. In our recent paper^9^, we developed a protocol called kTWAS (kernel-TWAS) using the ElasticNet model from PrediXcan to select variants and the well-known Sequence Kernel Association Test (SKAT) to associate variants to phenotype^21^. We demonstrated that kTWAS outperforms TWAS in real data and simulations under different genetic architectures^9^. Independently, another group at Emory University has released VC-TWAS (Variance Component TWAS)^10^, which utilizes a Bayesian linear model adapted from Tigar^8^ to select features and an equivalent kernel test for aggregation. Although these two publications use different approaches to model GReX and different parameterizations to conduct simulations, both studies conclude that kernel methods outperform GReX models in feature aggregation. In this work, we extend these findings by also replacing GReX-directed feature selection with a straightforward method for selecting variants based on their marginal effects on expression. By thoroughly comparing two GReX-based protocols and two protocols that disentangle feature selection from aggregation, we show that separating feature selection and aggregation into different statistical models significantly improves the power of TWAS in many conditions. This clearly shows that GReX models are not always optimal for either feature selection or aggregation, and future research should consider GReX as just one of several components that can be chosen to maximize the power of TWAS in a two-step framework.

Although TWAS is most often conducted on summary statistics (i.e., meta-analysis) rather than subject-level genotypes^1,3,7,22^, our previous results show that the relative power between protocols utilizing summary statistics is consistent with the relative power of their counterparts utilizing subject-level genotype data^9^. We therefore chose to analyze subject-level data in order to simplify the comparison between GReX-based and disentangled TWAS protocols. For the same reason, we also restricted our comparisons to cis-genetic elements and single-tissue analyses to avoid possible complications introduced by the integration of more advanced models.

## RESULTS

### Novel TWAS protocol disentangling feature selection and aggregation

We implement the novel protocol called Marginal + Kernel TWAS (abbreviated mkTWAS), which replaces GReX with marginal effect-based feature selection and kernel-based feature aggregation (implemented with FastQTL^23^ and SKAT^21^ respectively). For a given focal gene, we first select genetic variants by associating individual variants with the gene’s expression level^23^, retaining significant variants as potential expression quantitative trait loci (eQTLs) for downstream association mapping. We then aggregate potential eQTLs using SKAT’s kernel-based score test to determine gene to phenotype associations^21^ (**Online Methods**). We include a relatively large number of potential eQTLs (with nominal p-values less than 0.05 before multiple-test correction) as input to SKAT, since our previous work found that the performance of SKAT favors a large number of weakly correlated variants over a small number of highly significant variants. We hypothesize that this is because kernel methods are more robust to noise and therefore extract weaker signals, allowing statistical power to scale with an increasing number of features.

To evaluate the effectiveness of disentangled feature selection and aggregation, we compare a total of four protocols (**Table 1**). Protocols (1) and (2) use GReX to bind together feature selection and aggregation. Protocol (1), referred to as GReX (ElasticNet), adopts the ElasticNet linear model from PrediXcan^6^ to estimate GReX. Protocol (2), referred to as GReX (BSLMM), adopts the Bayesian sparse linear mixed model (BSLMM) to estimate GReX. As the existing BSLMM tool Fusion^7^ operates on summary statistics, we instead incorporate the weights of the BSLMM model into PrediXcan for association mapping. Protocols (3) and (4) separate feature selection and aggregation into different models. Protocol (3), referred to as ElasticNet + Kernel, uses the ElasticNet model from PrediXcan^6^ for feature selection and SKAT^24^ for feature aggregation as implemented in our previous method kTWAS^9^. Protocol (4), referred to as Marginal + Kernel, uses marginal genotype-expression effects for feature selection and SKAT for aggregation, as described above in our novel method mkTWAS. Type-I error is experimentally quantified by simulating under the null hypothesis for each protocol. Details of each protocol and the type-I error simulations are found in **Online Methods**.

**Table 1:** Design of compared protocols

| Protocol No. & Name | Feature Selection | Feature Aggregation | Implementation |
| --- | --- | --- | --- |
| 1) GReX (ElasticNet) | GReX: ElasticNet | GReX: ElasticNet | PrediXcan |
| 2) GReX (BSLMM) | GReX: Bayesian model | GReX: Bayesian model | PrediXcan/BSLMM |
| 3) ElasticNet + Kernel | GReX: ElasticNet | Kernel | PrediXcan + SKAT |
| 4) Marginal + Kernel | Marginal effects | Kernel | FastQTL + SKAT |

### Real data analysis

We first compare the four protocols above by analyzing WTCCC genotype data^25^, with complete outcomes listed in **Supplementary Tables S1 - S3**. For quantitative evaluation, we chose type 1 diabetes (T1D) and rheumatoid arthritis (RA) out of seven possible WTCCC diseases, as all four protocols discovered a large number of candidate genes in these diseases (p-value less than 0.05 after Bonferroni correction). For both diseases, Marginal + Kernel identifies the largest number of significant genes out of all protocols (**Supplementary Tables S1 and S2**). To validate the functional relevance of the identified genes, we refer to the DisGeNET database of human gene-disease associations^26,27^. We assess each protocol on the number of their discovered genes which are reported as disease-associated in DisGeNET (successes), as well as the proportion (success ratio) of these validated genes among all of the significant genes identified by the given protocol (**Supplementary Table S4**). Due to implementation differences, Marginal + Kernel and GReX (BSLMM) can assess all 19,696 genes in DisGeNET, whereas ElasticNet + Kernel and GReX (ElasticNet) only assess 7,252 genes for which corresponding ElasticNet models are available from the PrediXcan website^6^. As such, we only compare between the pairs Marginal + Kernel versus GReX (BSLMM) and ElasticNet + Kernel versus GReX (ElasticNet), giving a total of four comparisons over two diseases. In three of these four comparisons, the protocols disentangling feature selection and feature aggregation (Marginal + Kernel, ElasticNet + Kernel) outperform GReX-based protocols in both number and proportion of successfully identified genes (**Fig. 2 a,c**). The only exception is between Marginal + Kernel and GReX (BSLMM) in T1D, where Marginal + Kernel has a slightly lower success ratio but a much higher number of successes (**Fig. 2 a,c**). Since protocols which identify fewer genes tend to find a higher proportion of known disease-associated genes, we also compare the number and proportion of genes which are exclusively identified by only one of the two protocols under comparison (**Online Methods**). Complete outcomes are listed in **Supplementary Table S5**. Again, in three out of four comparisons the disentangled protocols outperform GReX-based protocols (**Fig. 2 b,d**). The only exception is between ElasticNet + Kernel and GReX (ElasticNet) in T1D, where ElasticNet + Kernel has a lower success ratio but identifies a much larger number genes (**Fig. 2 b,d**). We perform an additional comparison between the two disentangled protocols Marginal + Kernel and ElasticNet + Kernel to evaluate the effectiveness of GReX on feature selection alone. We find that the simple marginal effects model used in Marginal + Kernel outperforms the more complex ElasticNet model used in ElasticNet + Kernel in total number of successes, but has a lower success ratio (**Fig. 3 a,c**). However, when omitting genes identified by both protocols, Marginal + Kernel substantially outperforms ElasticNet + Kernel in both the total number and the ratio of exclusive successes (**Fig. 3 b,d**).

**Figure 2:**
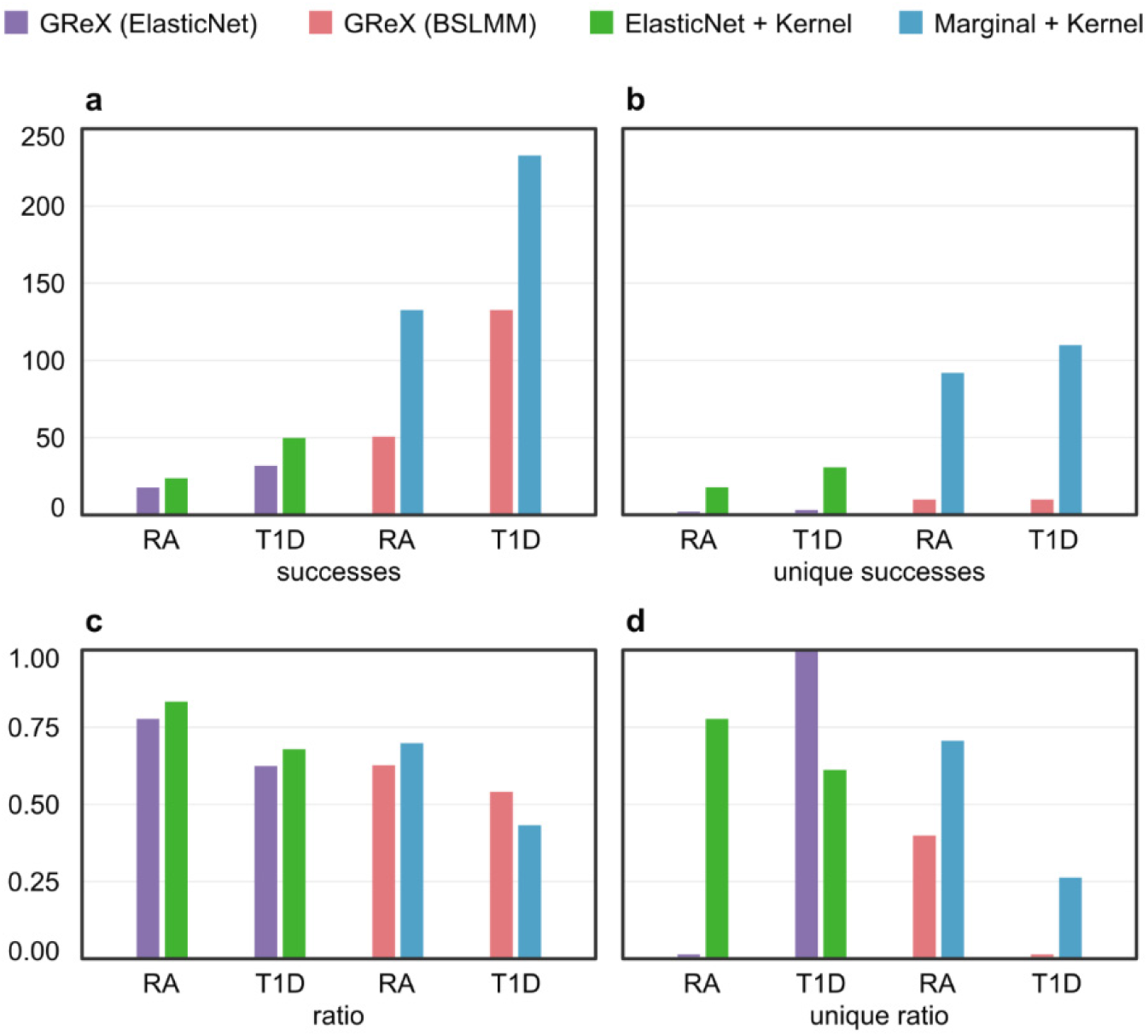
Comparison of GReX-based versus disentangled protocols in WTCCC data. Each figure compares two pairs of protocols in which the same number of genes are assessed over two WTCCC diseases (T1D and RA): GReX (ElasticNet) versus ElasticNet + Kernel (left), and GReX (BSLMM) versus Marginal + Kernel (right). **(a)** Total number of discovered genes (successes) which are reported as disease-associated in DisGeNET. **(b)** Number of validated genes (successes) discovered exclusively by one of the two protocols under comparison. **(c)** Proportion (success ratio) of all discovered genes which are validated by DisGeNET. **(d)** Proportion (success ratio) of genes discovered exclusively by each protocol which are validated by DisGeNET.

**Figure 3:**
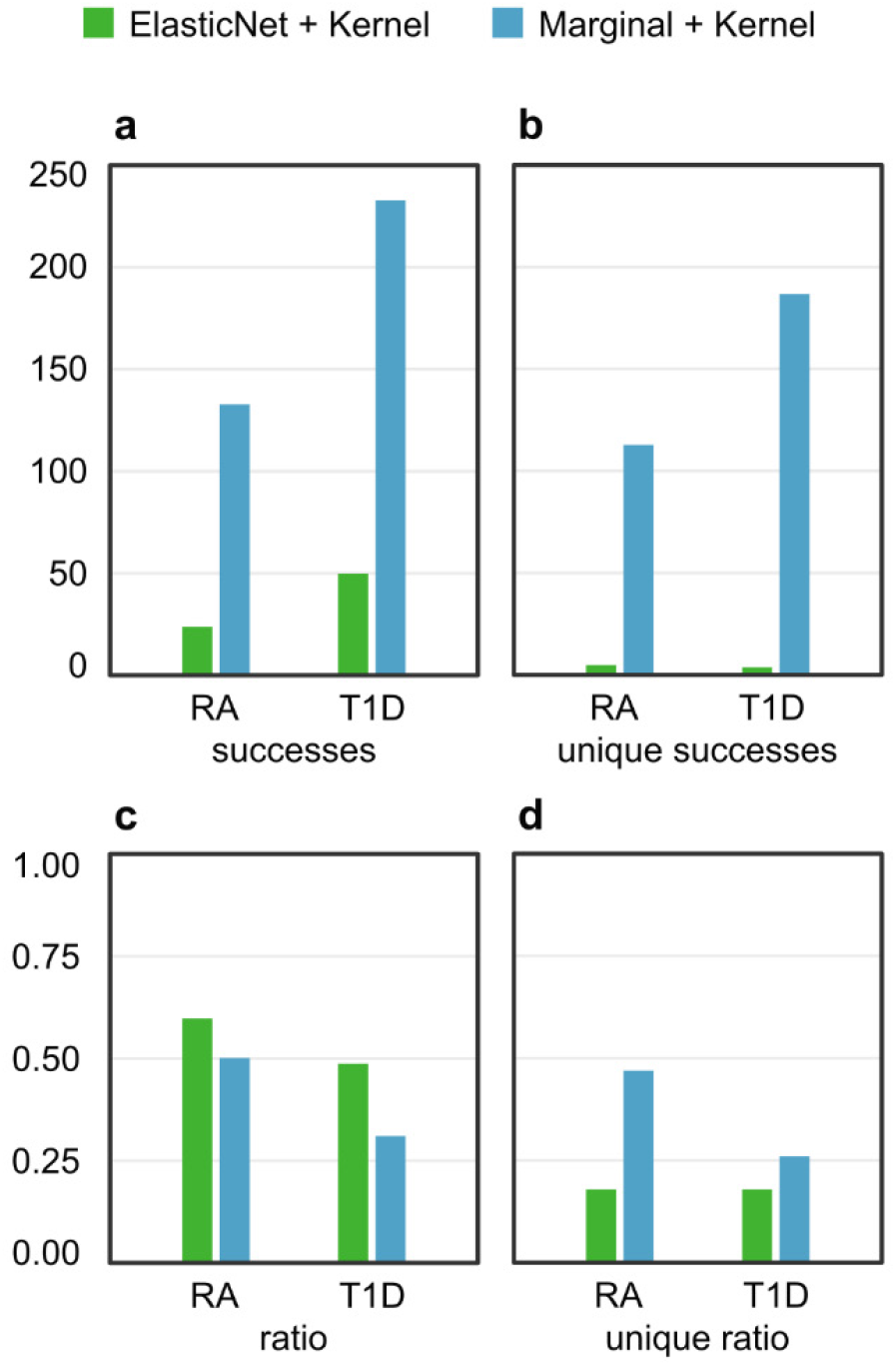
Comparison of marginal effect- and ElasticNet-based feature selection in WTCCC data. The two disentangled protocols ElasticNet + Kernel (left) and Marginal + Kernel (right) are compared on two WTCCC diseases (T1D and RA). Both protocols use kernel-based feature aggregation, but have different feature selection methods. **(a)** Number of discovered genes (successes) which are reported as disease-associated in DisGeNET. **(b)** Number of validated genes (successes) discovered exclusively by one of the two protocols under comparison. **(c)** Proportion (success ratio) of all discovered genes which are validated by DisGeNET. **(d)** Proportion (success ratio) of genes discovered exclusively by each protocol which are validated by DisGeNET.

In addition to the above comparisons, for each disease we investigate whether the protocols can identify the top three genes with the highest gene-disease association scores in DisGeNET. The DisGeNET score takes into account the number of sources that report an association, the type of curation for each source, animal models where the association was studied, and the number of supporting publications discovered via text mining. This evidence is combined to score each gene by the confidence of its gene-disease association. The top three genes for RA are TNF, PTPN22, and SLC22A4, of which Marginal + Kernel is able to detect TNF and PTPN22, whereas none of the other protocols can identify any of the three genes. For T1D, the top three genes are PTPN22, INS, and HNF1A, of which Marginal + Kernel identifies PTPN22, whereas the remaining protocols do not identify any of the three genes.

Following standard practice in methodological works^6–8,14,28^, we search the literature for additional evidence that the identified genes are relevant to disease. As discussed above, the only protocol which associates PTPN22 with T1D and RA is Marginal + Kernel. PTPN22 is highly scored in DisGeNET and has extensive literature support^29^. Among the five WTCCC diseases with an insufficient number of identifiable genes for quantitative comparison, Marginal + Kernel is the only protocol which associates TCF7L2 with type 2 diabetes and IRGM with Crohn’s disease. Both genes are well-supported by literature^30–33^. Based on our literature search, the top 5 significant genes for all four protocols are generally well supported. A comprehensive listing and discussion of the relevant literature for each gene is included in **Supplementary Notes** and **Supplementary Table S6**.

### Simulations

To thoroughly investigate the conditions under which disentangling feature selection and aggregation outperforms GReX alone, we conduct simulations comparing the four protocols in two scenarios: causality, where genotype causes phenotype via the intermediary of expression, and pleiotropy, where genotype causes phenotype and expression independently (**Supplementary Fig. S1**). As previous publications show that TWAS enjoys higher power in pleiotropy than causality^10,34,35^, the pleiotropic simulations have reduced heritability to better distinguish power differences between each protocol (**Online Methods**). In both scenarios, we simulate expression and phenotype with an additive genetic architecture and three interactive architectures labelled epistatic, compensatory, and heterogeneous (**Fig. 4 & 5**). Under pleiotropy, the two disentangled methods (Marginal + Kernel and ElasticNet + Kernel) significantly outperform the GReX-based protocols (**Fig. 4**). Marginal + Kernel also outperforms ElasticNet + Kernel, showing that a simple marginal effect-based model can outperform a regularized ElasticNet model in feature selection (**Fig. 4**). Under causality, the GReX-based protocols have higher power in the additive case, with GReX (BSLMM) leading (**Fig. 5 a,b,c**). This is unsurprising since the additive architecture consists of the same causal relations assumed by GReX (genotype causes expression and expression causes phenotype). In the interaction architectures under causality, all protocols have similar power with BSLMM leading slightly (**Fig. 5 d,e,f**). Detailed mathematical formulas and parameterizations of our simulations are available in **Online Methods**.

**Figure 4:**
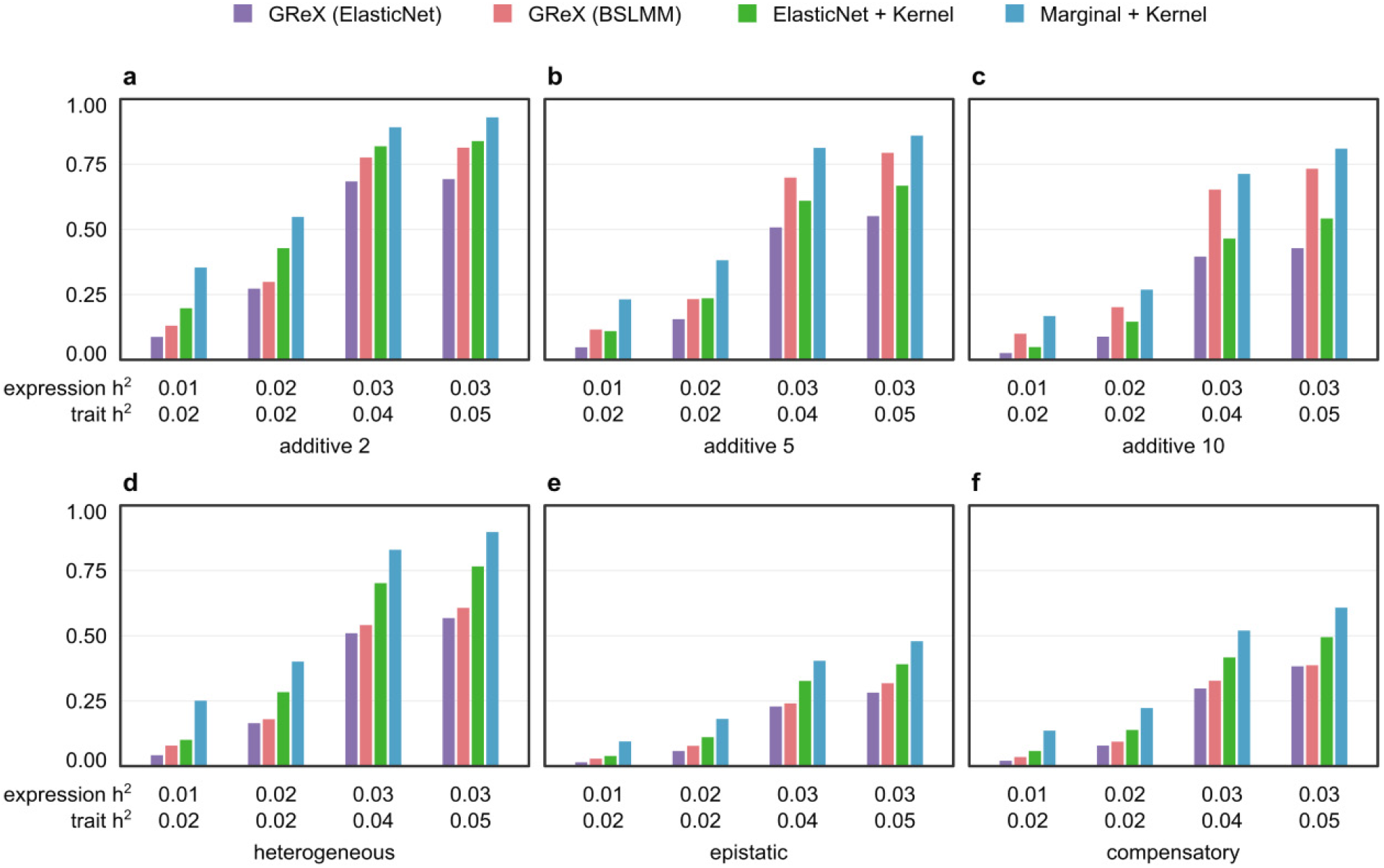
Power comparison of protocols in simulated pleiotropy scenario. The pleiotropic scenario simulates independent associations from genotype to phenotype and expressions (see **Supplementary Fig. S1** and **Online Methods**). Power is indicated on the y-axis. **(a) - (c)** are results under an additive genetic architecture, with differing expression heritability and local trait heritability denoted below each panel. The total number of contributing genetic variants is 2, 5, and 10 in each panel (left to right). **(d) - (f)** are results under interaction architectures, with expression heritability and local trait heritability denoted below. From left to right, the specific interactions are heterogeneous (logical ‘OR’), epistatic (logical ‘AND’), and compensatory (logical ‘XOR’).

**Figure 5:**
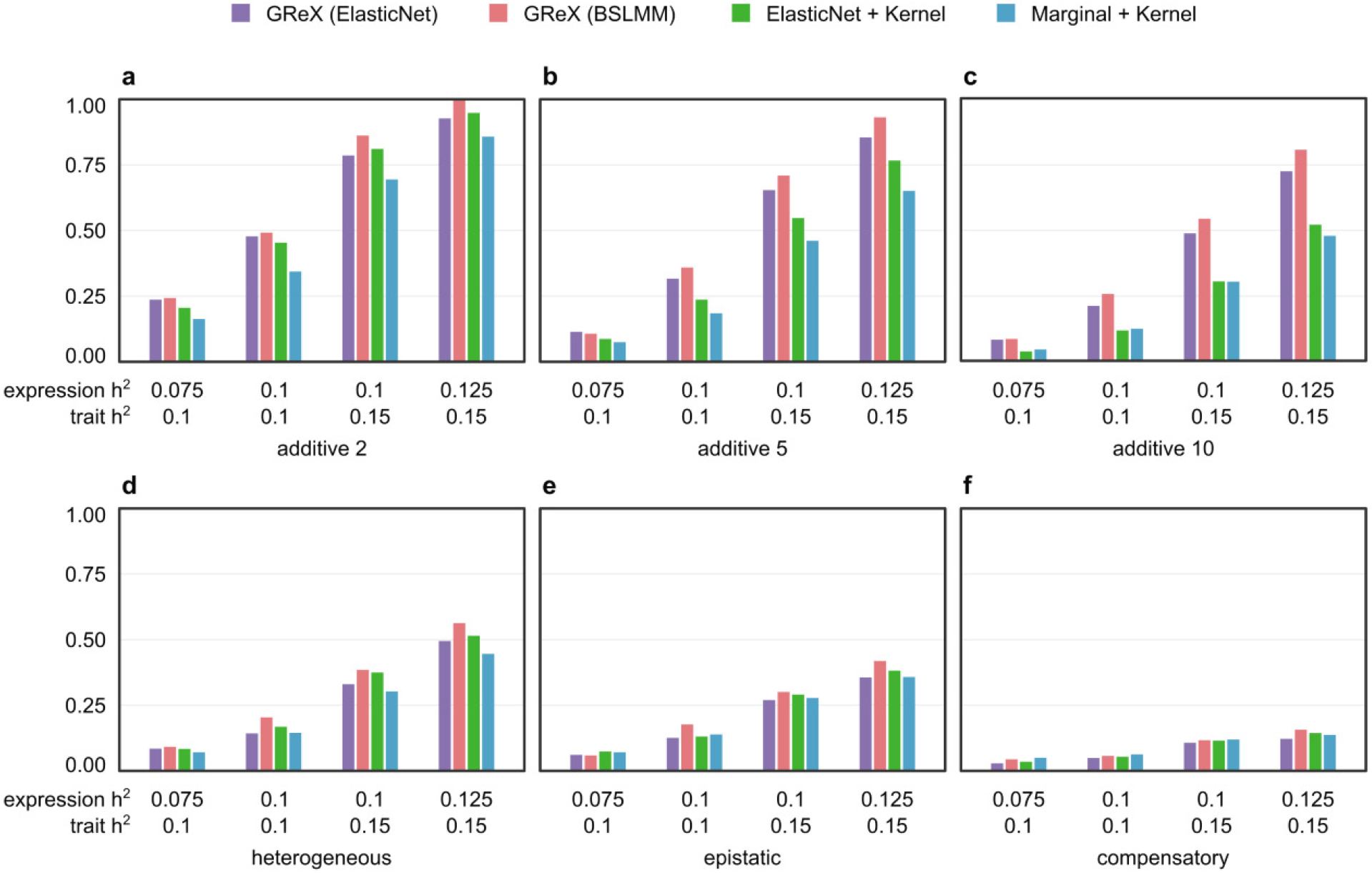
Power comparison of protocols in simulated causality scenario. The causality scenario simulates dependence of phenotype on genotype via gene expression (see **Supplementary Fig. S1** and **Online Methods**). Panels **(a) – (f)** have the same layout as figure 4.

## DISCUSSION

Our results show that in most cases, decoupling feature selection and aggregation allows even a simple feature selection method, based on the individual effects of genetic variants, to outperform a complex regularized model, which utilizes the combined linear effects of all potential eQTLs. Combined with the two preceding publications which apply kernel-based feature aggregation to TWAS^9,10^, this clearly demonstrates that GReX is not an optimal choice for all conditions. Although GReX has been a successful approach for leveraging expression data in GWAS, it is inherently limited by its use of a single linear model to solve two high-dimensional machine learning problems. Current TWAS development has largely treated GReX as a monolithic component, perhaps because the underlying statistical understanding of GReX as a genotype (not expression) model has been overlooked. By separating feature selection and feature aggregation into independent procedures, we show that many potential combinations of methods for conducting TWAS have been overlooked, some of which can yield improved power and specificity in commonly seen genetic architectures.

The simplicity of our marginal effects model suggests that feature selection plays an underappreciated role in TWAS and deserves further investigation. One interpretation of our marginal method’s effectiveness is that it is more lenient in selecting variants, which allows a wider number of genes with poorly predicted expressions to be analyzed. For instance, PrediXcan can only analyze around 1/3 of the genes which have well-predicted expressions, whereas Marginal + Kernel can analyze all available genes in the transcriptome. We also propose that the larger number of variants selected by marginal effects pairs better with kernel-based aggregation, due to the previously discussed robustness of the kernel test to noise. These findings suggest that while feature selection and aggregation methods can be independently developed, it is also necessary to consider their compatibility when integrated in a two-step TWAS framework.

In order to simplify the design and interpretation of the protocols in this study, we did not consider trans-eQTLs and multiple tissue-based methods. As a future work, we will examine whether the conclusions of this study remain valid when potential trans-eQTLs are included in the protocols and simulations. Our findings can also apply to other types of middle-omic directed association mapping studies such as PWAS on proteins^36,37^ and IWAS^38^ on brain images. This opens many additional opportunities for applying new or existing combinations of tools to various datasets.

## Supporting information

Supplementary Figure and Literature Support

## Acknowledgements

Q.L. is supported by an NSERC Discovery Grant (RGPIN-2017-04860), a Canada Foundation for Innovation JELF grant (36605), a New Frontiers in Research Fund (NFRFE-2018-00748), an HBI Pilot grant and an ACHRI Startup grant. C.C. is supported by an ACHRI scholarship. D.K. is supported by an NSERC USRA award.

## ONLINE METHODS

### Notations

In this section, we use ***X*** to denote a matrix of genotypes over *k* × *n* individuals and genetic variants, and ***x, y***, and ***z*** to denote vectors of genetic variants, phenotypes, and gene expressions respectively. We use ***β*** to denote vectors of coefficients for genetic markers and ***ε*** for residuals. Vectors corresponding to a particular variant site are indexed by the subscript *i*.

### Analytic protocols

Four protocols were applied in simulations and real data analysis, with two GReX-based protocols chosen to represent flagship TWAS methods in practical use. (1) uses ElasticNet^39,40^ implemented by PrediXcan^6^, as a representative of regularized GReX models, which is applied to both feature selection and aggregation. (2) uses BSLMM^13^ as a representative of Bayesian GReX models, also applied to both feature selection and aggregation. As the widely used BSLMM tool Fusion operates on summary statistics^7^, we instead use PrediXcan to conduct subject-level association mapping from the weights estimated by BSLMM.

Two additional protocols were chosen to represent methods separating feature selection and aggregation. (3) combines ElasticNet feature selection with Kernel-based feature aggregation. Expression data is used to train an ElasticNet model such that 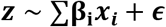, where the objective function minimizes 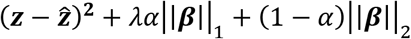. Training is conducted using the R package glmnet^39,40^ in simulations, while pre-trained coefficients from the PrediXcan website are used in real data analysis (http://predictdb.org). Unlike the standard TWAS protocol however, the predicted expressions are not used directly to conduct association mapping. Instead the weighted genetic variants are formed into a kernel ***K = X’DX**/n*, where ***X*** is a matrix of the selected variants, ***D*** is a diagonal matrix of variant weights, and *n* is the number of genetic variants. Using SKAT, we conduct a score-test ***Q = y’Ky***, where ***K*** is the kernel described above. Complete details are in our recent publication^9^, and the code is available on GitHub (https://github.com/theLongLab/kTWAS). (4) Marginal + Kernel: This protocol uses FastQTL^23^ to carry out eQTL analyses on each gene and select a large number of genetic variants from potential eQTLs (variants with a nominal p-value lower than 0.05, without multiple-test correction). Note that the marginal effect of each variant is computed individually from the eQTLs. Selected variants are formed into the same kernel and SKAT score test as described in protocol (3), except the diagonal matrix ***D*** contains the log-base-10 p-values from eQTL mapping. The code is available on GitHub (https://github.com/theLongLab/mkTWAS).

### Type-I error estimation

Although both GReX-based protocols (1) and (2) are well-established and the type-I error of protocol (3), ElasticNet + Kernel, was recently assessed^9^, the type-I error may still vary depending on the simulations and implementations. We therefore generated random phenotypes using individual data from the 1000 Genomes Project^41^ to measure the type-I error for each protocols. The type-I error is estimated using the top 5% cutoff for the most significant p-values obtained by the null hypothesis simulation. As shown in **Supplementary Table S7**, all protocols have type-I errors comparable to their theoretical values.

### Real data analysis

For all four protocols, feature selection was performed on GTEx whole blood data^42^. Association tests were conducted on genotype data for seven diseases in WTCCC^25^. Out of 393273 features (SNPs) recorded in WTCCC, 363217 are shared with GTEx and utilized for feature selection. The sample size for each of the seven WTCCC diseases are listed in **Supplementary Table S8**.

#### Success rate analysis

To assess the relevance of the genes identified by each protocol, we examine the proportion of discovered genes which have annotations in the DisGeNET database^26,27^. Specifically, for any pair of protocols *A* and *B* yielding corresponding sets of associated genes, we take the difference of the sets *A – B* (genes only in *A* and not *B*), and *B - A* (genes only in *B* but not A), and find the proportion of genes in each set difference which are annotated in DisGeNET. These two proportions define the success rates of each protocol as presented in **Figs. 2 and 3**.

### Simulations

Gene expressions are simulated using genotype information from GTEx^42^, and phenotypes are simulated using information from the 1000 Genomes Project^41^.

#### Causal scenarios

We simulate the two commonly assumed scenarios pleiotropy and causality (**Supplementary Fig. S1**). All variants are sampled from a region including the relevant gene body and 1Mb of flanking sequences at both sides. Under pleiotropy, the phenotype ***y*** and expression ***z*** are independently caused by the same genetic variants, so that ***z** = f(**X***) + ***ε*** and ***y*** = *g_p_*(***x***) + ***ε***. Under causality, phenotype is caused by genotype directly and also via the intermediary effect of expression, so that ***z*** = *f*(***X***) + ***ε*** and ***y*** = *g_c_*(***z, x***) + ***ε***. Note that the function *f* which maps genotype to simulated expressions is identical in both scenarios, but the functions *g_p_* and *g_c_* for simulating phenotype differ.

#### Genetic architecture models

The functions *f, g_p_* and *g_c_* are defined differently depending on the specific genetic architecture. In the additive model, given *n* genetic variants in a genotype matrix ***X*** where ***X*** = {*x*_1_, …, *x_n_*}, the expression model is defined as 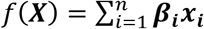. We set *n* as 2, 5, and 10 in our simulations. The effect size ***β_i_*** is drawn from the standard normal distribution *N*(0,1). In the interaction model, two genetic variants are chosen to affect gene expression or phenotype through one of three definitions. The ‘heterogeneous’ model is equivalent to the logical operation ‘OR’, in which the presence of a mutant allele in either or both variant sites causes a phenotypic change. The ‘epistatic’ model is equivalent to the logical operation ‘AND’, in which phenotypic change occurs only when a mutation is present at both variant sites. Finally, the ‘compensatory’ is equivalent to the logical operation ‘XOR’, in which a mutant allele can cause phenotypic change at either site, but if mutations occur at both sites their effect is negated. In all of the above models, the genetic component contributing to expression or phenotype is simulated as a value between 0 and 1, which is later rescaled based on expression or trait heritability. Under pleiotropy, *g_p_*(***X***) is defined identically to *f*(***X***), except that the variance component is rescaled by expression heritability instead of local trait heritability. Under causality, the additive genetic architecture defines *g_c_*(***z,x***) = ***z*** + ***ε***. In the interaction architectures, letting 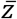 denote the median of the gene expression ***z*** we define

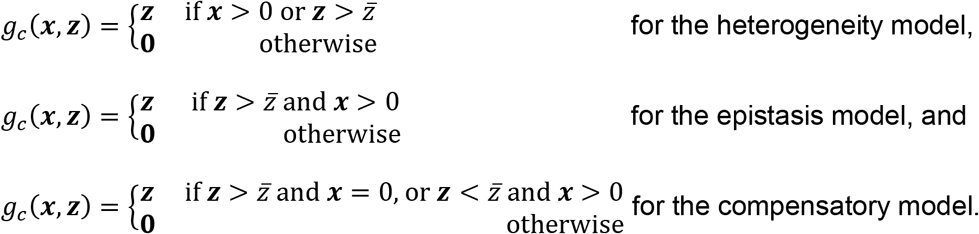

Illustrated examples on the evaluation of the above formulas are provided in **Supplementary Table S9**.

#### Variance component

The residual ***ε*** is randomly drawn from a normal distribution *N*(0, *σ*^2^) where the parameter *σ*^2^ is a scaling parameter which ensures that the expression heritability or trait heritability, denoted *h*^2^, is maintained at a pre-specified value. Specifically, let *G* denote the genetic component of expression or phenotype which is calculated from *f*(***X***), *g_p_*(***z***), or *g_c_*(***z,x***). Then *σ*^2^ is derived using the equation (*Var*(*G*) + *σ*^2^)/*σ*^2^ = *h*^2^, where *h*^2^ is pre-specified for a particular simulation. This ensures that the simulated expressions or phenotypes have the desired level of heritability.

### Web Resources

mkTWAS, https://github.com/theLongLab/mkTWAS

kTWAS https://github.com/theLongLab/kTWAS

BSLMM http://www.xzlab.org/software.html

PrediXcan, https://github.com/hakyim/PrediXcan

1000 Genomes Project, https://www.internationalgenome.org/

GTEx, https://gtexportal.org/

WTCCC, https://www.wtccc.org.uk/

---

**Supp Table S1: All significant genes identified by any of the four protocols in RA**. Significant ones are noted in bold. Note that different protocols have different number of Bonferroni corrections, as noted under the name of the protocols.

**Supp Table S2: All significant genes identified by any of the four protocols in T1D**. Significant ones are noted in bold. Note that different protocols have different number of Bonferroni corrections, as noted under the name of the protocols.

**Supp Table S3: All significant genes identified by any of the four protocols in other WTCCC diseases, i.e., CD, BD, HT, CAD, and T2D**. Significant ones are noted in bold. Note that different protocols have different number of Bonferroni corrections, as noted under the name of the protocols.

**Supp Table S4: Numbers and percentages of genes that are successfully annotated for all protocols**.

**Supp Table S5: Numbers and percentages of uniquely discovered genes that are successfully annotated for all protocols**.

**Supp Table S6: Literature support for a subset of the discovered genes**.

**Supp Table S7: Estimated empirical type-I errors of all protocols**.

**Supp Table S8: Case size and control size for all WTCCC diseases**.

**Supp Table S9: An interaction table for expressions and genotypes**. Assuming the expression ranges from 0.0 to 1.0, with 0.5 as its median, the tables below show how the interactions are defined.

## Notes

### Competing Interest Statement

The authors have declared no competing interest.

