## Supplementary Figure and Literature Support for "Disentangling genetic feature selection and aggregation in transcriptome-wide association studies"

**Supplementary Figure S1: Pleiotropy v.s. casualty.** Two commonly assumed scenarios are simulated. **(a)** Under causality, the genotype $x$ is causal to gene expression $z$, and in turn expression is causal to phenotype $y$, resulting in dependence of $y$ on both $z$ and $x$. (b) Under pleiotropy, the genotype $x$ is independently causal to phenotype $y$ and expression $z$. As such, the phenotype and expression are not causal to each other.

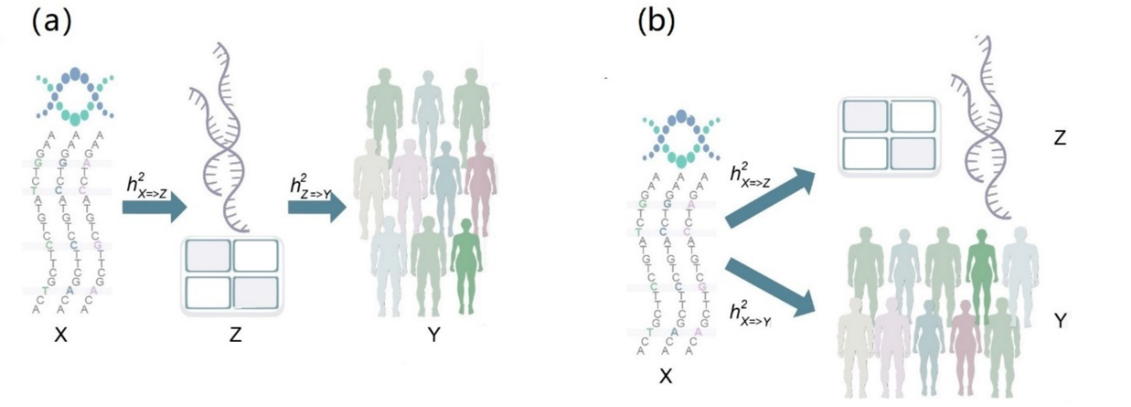

**Literature support for discovered genes**

For rheumatoid arthritis (RA), the majority of significant genes detected by Marginal + Kernel are within the Major Histocompatibility Complex (MHC) region. The MHC shows associations with almost all known autoimmune diseases as well as many inflammatory and infectious diseases[1]. Of the five most significant genes discovered by Marginal + Kernel (HLA-DRB1, LY6G5B, HLA-DMA, FKBPL and HLA-DQB1-AS1), three of them are from the human leukocyte antigen (HLA) gene family, all of which are located in the MHC. Genes from the HLA region have been reported as the most powerful disease risk predictors for RA[2]. In addition, there exist significant differences in the expression of FKBPL in some tissues for rheumatoid arthritis[3]. The majority of the five most significant genes reported by the other three protocols under comparison are also supported by existing literature (**Supplementary Table S6**).

For type 1 diabetes (T1D), the majority of significant genes detected by Marginal + Kernel are also located in the MHC region. Of the five genes reported by Marginal + Kernel as most strongly associated with T1D (TAP1, HLA-DQA1, CFB, HLA-DQB1, and HLA-DRB5), three are from the HLA region (HLA-DQA1, HLA-DQB1, and HLA-DRB5). The HLA region accounts for approximately half of the familial aggregation of T1D[4]. In particular, polymorphisms of class II HLA genes encoding DQ, DR, and to a lesser extent DP are the major genetic determinants of T1D[4]. HLA-DQA1, HLA-DQB1, and HLA-DRB5 are all subunits of DQ and DR. TAP1, which Marginal + Kernel reports as the most significant gene polymorphism, has evidence of association with T1D[5]. The majority of the five most significant genes reported by the other three protocols in this study are also supported by previous literature (**Supplementary Table S6**).

A limited number of significant genes were identified for the five remaining diseases in WTCCC: bipolar disorder (BD), coronary artery disease (CAD), Crohn's disease (CD), type 2 diabetes (T2D), and hypertension (HT) (**Supplementary Table S3**). For example, transcription factor 7-like 2 (TCF7L2) is the most important susceptibility gene for type 2 diabetes identified by Marginal + Kernel[6]. Among the four protocols, only Marginal + Kernel was able to identify TCF7L2. Another example is the gene IRGM, which plays an important role in the pathogenesis of Crohn's disease and is recognized as an independent major CD susceptibility locus from several previous studies[7,8]. Marginal + Kernel is the only method which detected IRGM as significantly associated with CD. Supporting literature PMIDs are listed in **Supplementary Table S6**.
